# Poles apart: the structure and composition of the bird community in bamboo in the Eastern Himalaya

**DOI:** 10.1101/2023.01.27.525938

**Authors:** Sidharth Srinivasan, Aman Biswakarma, D.K. Pradhan, Shambu Rai, Umesh Srinivasan

## Abstract

Bamboos (Poaceae: Bambuseae) are a unique and diverse group of plants (> 1500 species) that are found in tropical and subtropical regions of the world. Various birds, mammals, and insects use bamboo either directly or indirectly, and some species are obligate bamboo specialists. For instance, several species of Neotropical birds are found in bamboo alone. However, the mechanisms underlying dependence on/associations with bamboo are almost entirely unknown. We studied bird communities in bamboo and rainforest across two seasons in the Eastern Himalaya, where we explored a possible mechanism for bird associations with bamboo – dietary resource specialization. We calculated use of bamboo versus rainforest by bird species and described the structure and composition of the bird and arthropod communities in both habitats and seasons. We provide the first systematic evidence of bamboo specialist bird species from the Eastern Himalaya. Although the species richness of birds and arthropods in both habitats and seasons did not vary starkly, we found that the composition of bird and arthropod communities in both habitats and both seasons to be distinct. Interestingly, we found arthropod communities in different substrates of bamboo to also be distinct. Bird specialization in bamboo in the Eastern Himalaya could be because of their dietary specialization to the unique arthropods found in bamboo. The results from this study emphasise the importance of bamboo in the Eastern Himalaya and provide baseline information that might aid in conservation.

## Introduction

Bamboos (Poaceae: Bambuseae) are distinct, diverse groups of plants that have intrigued scientists and naturalists due to their unique ecology. Bamboo is known as a ‘boom and bust’ resource due to its semelparous nature - most species are known to quickly colonise treefall gaps and landslides to become the dominant vegetation, then undergo a long vegetative growth phase, followed by a single reproductive event where all or most individuals flower and seed synchronously (mast flowering/seeding), and then die en-masse. The mast seeding events occur anywhere between three and over a hundred years depending on the species of bamboo (Janzen 1976). This makes bamboo a successional and an ephemeral habitat, with mature forests replacing it following mast seeding and hence making them ‘temporally unstable’ (Ramakrishnan 1992, Raman et al. 1998, Socolar et al. 2013). Bamboo is found in tropical and subtropical habitats around the world, with Asia harbouring the maximum species richness of bamboo, followed by South America (with less than half the species richness of Asia; (Bystriakova 2003, Bystriakova et al. 2004).

Numerous birds, mammals, and insects use bamboo either directly or indirectly (Dierenfeld et al. 1982, Schellerich-Kaaden et al. 1997, Wright et al. 2008, Areta et al. 2009 p. 2, Leite et al. 2013). Although many species inhabit bamboo and although bamboo is generally abundant in the areas where it is found, studies exploring the role of bamboo in shaping bird communities are few (Raman et al. 1998, Rother et al. 2013). Most of our knowledge of bamboo-bird relationships comes from the Neotropics, where over 90 species of birds across diverse families are being partial to bamboo (Areta et al. 2009). At least 20 species of birds are obligate specialists (species that are characterised by their occurrence in a particular habitat and their absence in others) on the bamboo genus *Guadua*, highlighting the importance of specific genera of bamboo (Areta et al. 2009). Most of these birds are insectivores, but three species of granivorous birds are unexpectedly dependent on seeds from the inherently ephemeral and rare bamboo mast-seeding events (Areta et al. 2009, Lees et al. 2021). Most birds specialising in Neotropical bamboo habitats are non-migratory insectivores that generally forage along the culms (bamboo stems) or within the dense foliage (Pierpont and Fitzpatrick 1983). These foraging behaviours are known to be unique, such as probing into the stems or the internodes of bamboo or ripping apart the smaller stems of bamboo (Hilty et al. 1979, Kratter 1995, Leite et al. 2013). These specialised behaviours might contribute to their greater abundances in bamboo stands when compared with the abundance of other species in more generalised habitats (Kratter 1997). Bamboo species also have small range sizes and therefore exist at higher population densities in bamboo stands (Kratter 1995). Certain specialist species only inhabit larger bamboo patches, with smaller patches harbouring a subset of these species (Lebbin 2013). Arthropod prey are also richer in bamboo, and the combination of the distinct vegetation structure and the richer arthropod resources might facilitate specialisation on bamboo in the Neotropics (Reid et al. 2004, Lebbin 2007). In other words, birds can specialise on the distinctive prey in specific substrates in bamboo or on the distinct structure of bamboo (perches provided by bamboo can be used by flycatchers, for example; Kratter 1995). Bamboo could also be important for generalist insectivorous species because of the richer arthropod community and/or increased refuge from predators arising from the higher cover provided by the dense foliage of bamboo (Reid et al. 2004, Rother et al. 2013, Socolar et al. 2013). This could contribute to the overall bird species richness, which is often higher in bamboo than in non-bamboo habitats and other monodominant habitats (Rother et al. 2013).

Bamboo has largely been ignored in systematic community-wide avian studies (Raman et al. 1998, Rother et al. 2013). This is especially true in South Asia, where bamboo species richness peaks. Bamboo has been studied as a land use/land cover class in a handful of studies from this region, especially from North-East India and the Indian Eastern Himalaya, but usually as a successional stage in forests recovering from slash-and-burn agriculture (*jhum*) and selective logging forests (both see a profusion of bamboo; Raman et al. 1998, Raman 2001, Borah et al. 2018, Srinivasan 2019). Bamboo might buffer the impacts of logging on birds; the survival rates of some insectivorous birds showed little or no change in logged forests (Srinivasan and Wilcove 2021). Anecdotal evidence suggests that a few bird species might be bamboo specialists in the Eastern Himalaya (Ali and Ripley 1971, Olson 1986) but systematic studies quantifying the degree to which birds specialise in bamboo and the mechanisms underlying such specialisation remains unknown.

Bamboo habitats harbour a richer community of arthropods in terms of their density and abundance in the Neotropics (Reid et al. 2004, Lebbin 2007). Globally, a number of arthropod species specialise on bamboo (Schellerich-Kaaden et al. 1997, Viraktamath and Webb 2019), potentially altering the composition of the arthropod community in bamboo when compared with forests. Several arthropod groups also specialise on different substrates in bamboo – aquatic and semi-aquatic insects, especially mosquito larvae (Louton et al. 1996, Campos 2013) and bugs (Kovac 2000) in the internodes, bugs and beetles on the leaves and culms sheaths (coverings on the stems of bamboo; Kovac 2000), and fruit flies on fresh bamboo shoots (Hancock and Drew 1999). In the Himalaya, arthropod abundance is the limiting resource for insectivorous birds, and songbird richness peaked where arthropod abundance was the highest (Ghosh-Harihar 2013, Supriya et al. 2020). These studies were restricted to forests, however, and such relationships in bamboo stands are yet to be explored.

Seasonality also plays an important role in determining the abundance, richness and community composition of both arthropods and birds. Seasonality affects the richness of birds in the Himalaya, with migrations occurring within and from outside the Himalaya (Somveille et al. 2013). Arthropods peak in abundance in summer, and winter is considered a period of resource scarcity for birds (Ghosh et al. 2011, Srinivasan et al. 2018). This apparent winter dip in arthropod abundance and its subsequent effect on the bird community has not been quantified in bamboo. The impacts of seasonal resource availability on bird communities in bamboo habitats is a lacuna in our understanding of avian assemblages in this region, especially because (apart from mature stands) bamboo is an important successional habitat following forest loss/degradation.

We carried out this study to further our understanding of bamboo-bird associations in the Eastern Himalaya. The main objectives of our study were:

1. To determine and compare the composition of bird communities in bamboo stands and rainforest.
2. To quantify resource availability for insectivorous birds and explore the possibility that dietary specialisation acts as a mechanism for differences in bird communities between the two habitats.

## Methods

### Study Area

We carried out this study in the Eastern Himalaya, a Global Biodiversity Hotspot (Myers et al. 2000). Bird species richness in the Himalaya is amongst the highest in the world (Orme et al. 2005) and within the Himalaya, richness peaks in the Eastern Himalaya, which harbours more than 650 species of breeding birds (Singh 1994, Srinivasan et al. 2010). Bamboo species richness in this region is also one of the highest in the world, with over 60 species being found in the Eastern Himalaya (Bystriakova 2003).

We carried out the study in the Eaglenest Wildlife Sanctuary (hereafter EWS; 27.03° to 27.15° N and 92.30° to 92.58° E), located in the West Kameng district in western Arunachal Pradesh. EWS harbours an extensive elevational gradient and receives an annual rainfall of over 2000mm, especially in the lower elevations, mostly received during the South-West monsoon (June to October; Price et al. 2011). The resulting vegetation types are diverse – from alpine and temperate forests at higher altitudes to montane and subtropical forests at middle elevations and tropical forests at lower altitudes.

We sampled bamboo and old-growth rainforest at altitudes ranging from 800-1200m (Figure 1). The vegetation in this area is tropical/sub-tropical wet evergreen forests (Champion and Seth 1968) with extensive mature bamboo stands. Bamboo grows on steep slopes in this region, covering entire hillsides. We selected two such hillsides, with the dominant bamboo species being *Dendrocalamus longispathus* and negligible patches of a non-sympodial (runny) bamboo species which was not identified. *Dendrocalamus longispathus* is a large, sympodial (clump-forming) bamboo with huge culms and canopies reaching roughly 20m in height. In the study area, this species forms near-monotypic bamboo-dominated patches covering over 30 ha., with a few very large trees interspersed within the habitat. We selected undisturbed primary forest patches as the rainforest plots. The sampling period for this study spanned two seasons: winter from mid-January to mid-March and summer, between mid-March and mid-May.

**Figure 1.**
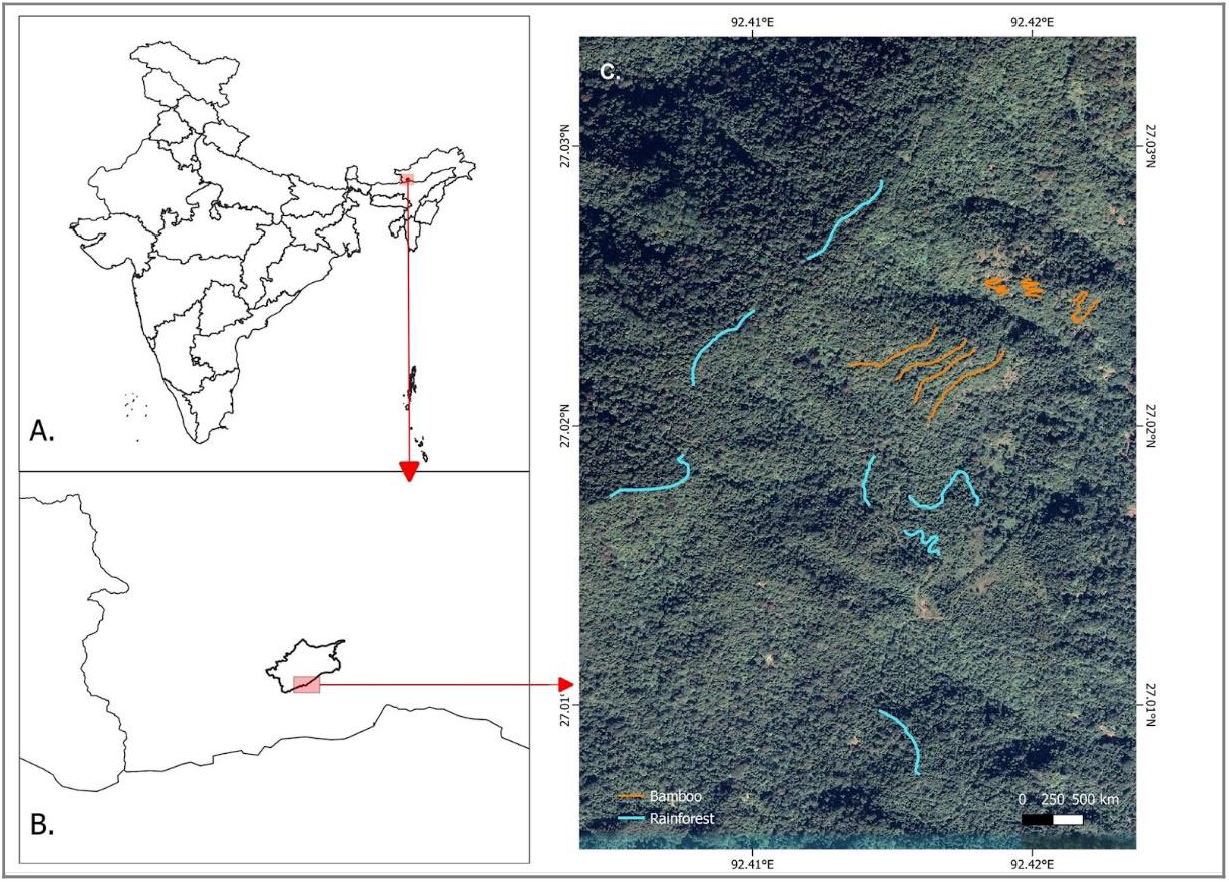
Map of India (A) showing the location of Eaglenest Wildlife Sanctuary (B) within north-east India. Transects in rainforest and bamboo are marked in blue and orange respectively (**C**).

## Sampling Design

### Bird Sampling

We sampled birds using line transects to estimate habitat use. We established two plots of similar sizes (∼30 ha) in each habitat – bamboo and rainforest. Within them, we sampled birds by establishing seven line transects in each habitat of varying lengths, from 200 to 500m. Because of the highly undulating terrain and the resulting inaccessibility, especially in the bamboo stands, adjacent transects are not likely to be independent of each other. We walked each transect at least 15 times in total and at least five times in each season, tallying a total of 386 walks of 14 transects while ensuring that the effort was similar across habitats in each season (47 vs 48.5 km in bamboo and rainforest in winter and 27 vs 24 km in bamboo and rainforest in summer). we walked transects between 0600 and 1100 hours, and also between 1400 and 1700 hours, depending on daylight hours in each season.

We detected and recorded birds either visually or aurally within a distance of 20m on either side of the transect, noting down species identity, group size (number of individuals) and the method of detection for each bird record. For gregarious species, an accurate group size measurement was not always possible; we therefore recorded group size in a range, from the minimum to the maximum number of individuals likely in the group. We used the midpoint of this range in subsequent analyses.

### Arthropod Sampling

To quantify resource availability, we sampled arthropods in both habitats. We selected 30 points at random in each habitat with a minimum distance of 30m between points. The points were randomly distributed around the transects, across each habitat. We sampled arthropods in these points by employing two methods – pitfall traps and branch beating.

Pitfall traps are an inexpensive and widely used technique to sample and estimate the relative abundances of ground-dwelling arthropods (Hohbein and Conway 2018), and provide an unbiased snapshot of the arthropod community (Brown and Matthews 2016). We placed open plastic containers (for 48 hours) with a diameter of 5 cm in the ground level with the soil surface. We filled the containers with a solution of detergent and water, roughly mixed in a ratio of 10:90.

Branch beating is another commonly used technique to sample foliage-dwelling arthropods. At each point, we selected branches randomly between 1-4 m above the ground, and placed them in a beating bag (a long funnel-shaped cloth bag), to which we attached a plastic container at the apex of the funnel. Branches were shaken and beaten for roughly 10 seconds to allow arthropods to fall into the bag and the plastic container, after which the branches were removed, and the container was detached and closed immediately.

We also sampled arthropods from within the culm sheaths (coverings on the bamboo stems) in bamboo by adapting the branch beating technique. We held the beating bag below the culm sheaths, as close as possible to the culms and we displaced the sheaths from the stems into the bag. we then shook the bag vigorously. We attached a plastic container to the smaller end of the bag which we used to collect culm arthropods. We could not replicate this procedure in summer, as the culm sheaths fall off bamboo stems after winter.

We sampled the same points in both seasons, totalling 270 points across both habitats (150 in winter and 120 in summer). We sampled only on relatively clear and sunny days, between 1000 and 1400 hours. We anaesthetized the collected arthropods using a cotton ball soaked in chloroform (Ghosh-Harihar 2013) and sorted and labelled them in the field. We then identified arthropods up to the order level.

## Analysis

### Habitat Use

For each bird species and in each season, we calculated a habitat-use metric by dividing the number of times it was detected in bamboo by the overall number of times it was detected in both habitats. We plotted these scores by season and arranged them in descending order of the scores. This metric is thus scaled from 0 to 1, where higher values indicate greater bamboo use and lower values indicate higher rainforest use. we defined bamboo specialists as those species with values to close or equal to one and rainforest specialists as those with score equal to or close to 0. We defined those species that seem to use both habitats equally, i.e., with a score of around 0.5, as generalists. we excluded rare species (species detected less than five times in each season and habitat) from this analysis.

### Bird Community Composition

We estimated bird species richness in each habitat and season using the jackknife procedure. To quantify differences in the composition of bird communities between bamboo and rainforest in summer and winter, we calculated a dissimilarity matrix using a site-by-species matrix based on raw counts of species in each habitat in each season. We defined a site as a transect in a particular season and habitat. We constructed the dissimilarity matrix by transforming the data using Hellinger distance, a dissimilarity measure that assigns lower weights to rare species (low non-zero counts), thereby reducing their influence on the overall dissimilarity (Buttigieg and Ramette 2014). We used a Non-Metric Dimensional Scaling (NMDS) to visualise the similarities/dissimilarities in habitat and seasonal bird community composition, by using the Bray-Curtis index on the transformed data (Bray and Curtis 1957).

To test whether the bird communities were significantly different between habitat and season, we used PERMANOVA (Permutational Analysis of Variance) with the Bray-Curtis index. We calculated the similarity percentage scores for species in each habitat and each season to determine the average contribution of individual species to the overall dissimilarity of bird communities between habitats. We then selected the species that were cumulatively contributing to 50% dissimilarity in each habitat and season (bamboo vs rainforest in winter and bamboo vs rainforest in summer). To understand species’ importance in influencing differences in community composition across habitats and seasons, we plotted these along with the habitat-use scores.

### Arthropod Community Composition

We analysed per-point arthropod abundance after removing the orders Hymenoptera and Isoptera. Hymenopteran abundances (mostly ants) increased considerably in the summer (especially in bamboo), making it harder to compare abundances across both seasons and habitats. Further, though we know that ants constitute the diets of some bird species, not all insectivorous birds are known to include ants as a part of their diet. Information on diet does not exist for bird species in this region, other than the fact that they are insectivorous. Ants are also known to compete with birds for the same arthropod prey, possibly excluding each other in areas where they are found (Supriya et al. 2020). Hence, we removed Hymentoptera before analyses. We encountered a single point in the bamboo pitfall traps in summer that harboured a hyper-abundance of Isoptera (termites; 875 individuals). Because this was clearly an outlier (as Isopteran abundances were generally much lower in other seasons and habitats), we also excluded this point from this analysis.

We calculated order level richness estimates for the arthropod communities in both habitats and seasons. To compare community composition of arthropods across habitats and seasons, we took an approach akin to the bird community composition analysis. We created a site-by-order matrix generated based on the raw abundances of orders in each habitat and season using the 30 random points as sites. We transformed the data using Hellinger distance to create a matrix of dissimilarity scores. To visualise the similarity in the community, we used an NMDS by using the Bray-Curtis index on the transformed data. We excluded rare orders (orders detected less than 3 times) and the data from the bamboo culm sheaths (as we could only sample them in winter) prior to this analysis.

During sampling in winter, we often observed birds seeking arthropods from the crevices of unopened bamboo culm sheaths. We hypothesised that the arthropod community in the sheaths were dissimilar from the arthropod community in the leaves. To test this hypothesis, we constructed a dissimilarity matrix of the arthropod community within the sheaths and from the branch beating procedure. After using a Hellinger distance to transform the data, we used the Bray-Curtis index to visualise the community similarity using an NMDS plot. We tested for the significance of dissimilarity in the arthropod communities in both the NMDS procedures using the PERMANOVA procedure with the Bray-Curtis index.

We used the *SPECIES* package (Wang 2011) to calculate species richness estimates and the package *vegan* (Oksanen et al. 2022) to perform the NMDS, PERMANOVA and similarity percentage scores analyses. We performed all analyses in the program R (R Core Team 2022).

## Results

### Species richness patterns and habitat use by birds

We recorded 5,337 observations of 186 species across rainforest and bamboo in both seasons. Bamboo yielded 2,160 observations of 160 species and rainforest 3,177 observations of 147 species. While the number of species between both seasons and between both habitats was similar (122 and 121 respectively), only 77 species were common across both habitats and seasons.

Estimated jackknife species richness was higher for rainforest than bamboo, irrespective of the season (155 vs 137 in winter and 150 vs 136 in summer, although 95% CIs overlapped between all estimates; Figure 2). Estimated richness declined marginally from winter to summer in both habitats.

**Figure 2.**
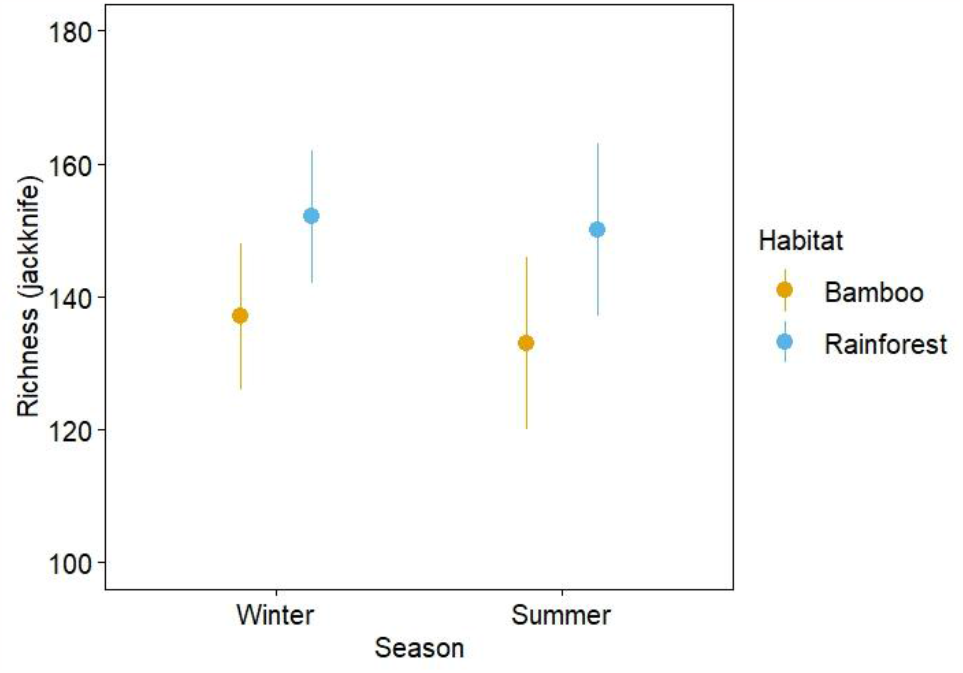
Estimated jackknife species richness of birds across habitats and seasons. Error bars represent 95% confidence intervals.

Generalists and specialists separated well when arranged from the highest to the lowest values of the habitat use score (where higher scores indicated more bamboo use and lower scores indicated more forest use). The species with the highest scores (one or close to one) in winter were the Collared Treepie (*Dendrocitta frontalis*), Pale-billed Parrotbill (*Chleuasicus atrosuperciliaris)*, Pale-headed Woodpecker (*Gecinulus grantia)*, Red-billed Scimitar Babbler (*Pomatorhinus ochraceiceps*), White-breasted Parrotbill (*Psittiparus ruficeps)*, Yellow-bellied Warbler (*Abroscopus superciliaris)* and the White-hooded Babbler (*Gampsorhynchus rufulus)*. These species also had the highest scores in summer, in addition to the Large Blue Flycatcher. we classify these species as ‘bamboo specialists’ (Figure 3).

**Figure 3.**
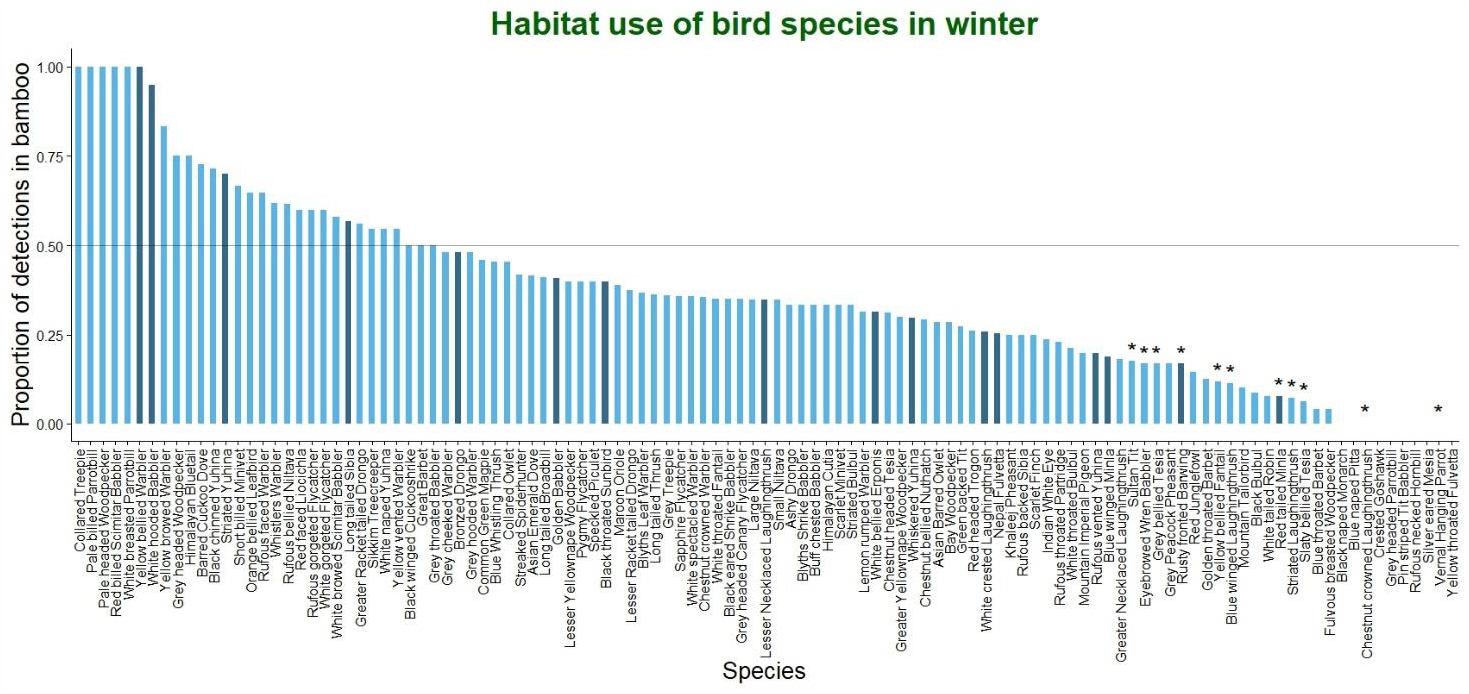
Habitat use scores of bird species in winter. Higher values indicate disproportionally higher presence in bamboo and lower scores indicate disproportionally higher presence in rainforest. Species are arranged in descending order of these values. Darker bars represent species that contribute to a cumulative 50% dissimilarity between the two habitats. Asterisk (*) above the bars indicates a significant contribution to the observed dissimilarity.

The Yellow-throated Fulvetta (*Schoeniparus cinereus*), Vernal Hanging Parrot (*Loriculus vernalis*), Pin-striped Tit-Babbler (*Mixornis gularis*), Rufous-necked Hornbill (*Aceros nipalensis*), Grey-headed Parrotbill (*Psittiparus gularis*), Chestnut-crowned Laughingthrush (*Trochalopteron erythrocephalum*) and the Blue-naped Pitta (*Hydrornis nipalensis*) had the lowest scores (zero or close to zero) in winter. In summer, in addition to the Yellow-throated Fulvetta and Pin-striped Tit Babbler, the Yellow-bellied Fantail (*Chelidorhynx hypoxanthus*), Rufous-gorgeted Flycatcher (*Ficedula strophiata*), Red Junglefowl (*Gallus gallus*) and Dark-sided Flycatcher (*Muscicapa sibirica*) also had the lowest scores. We consider these species rainforest specialists (Figure 4).

**Figure 4.**
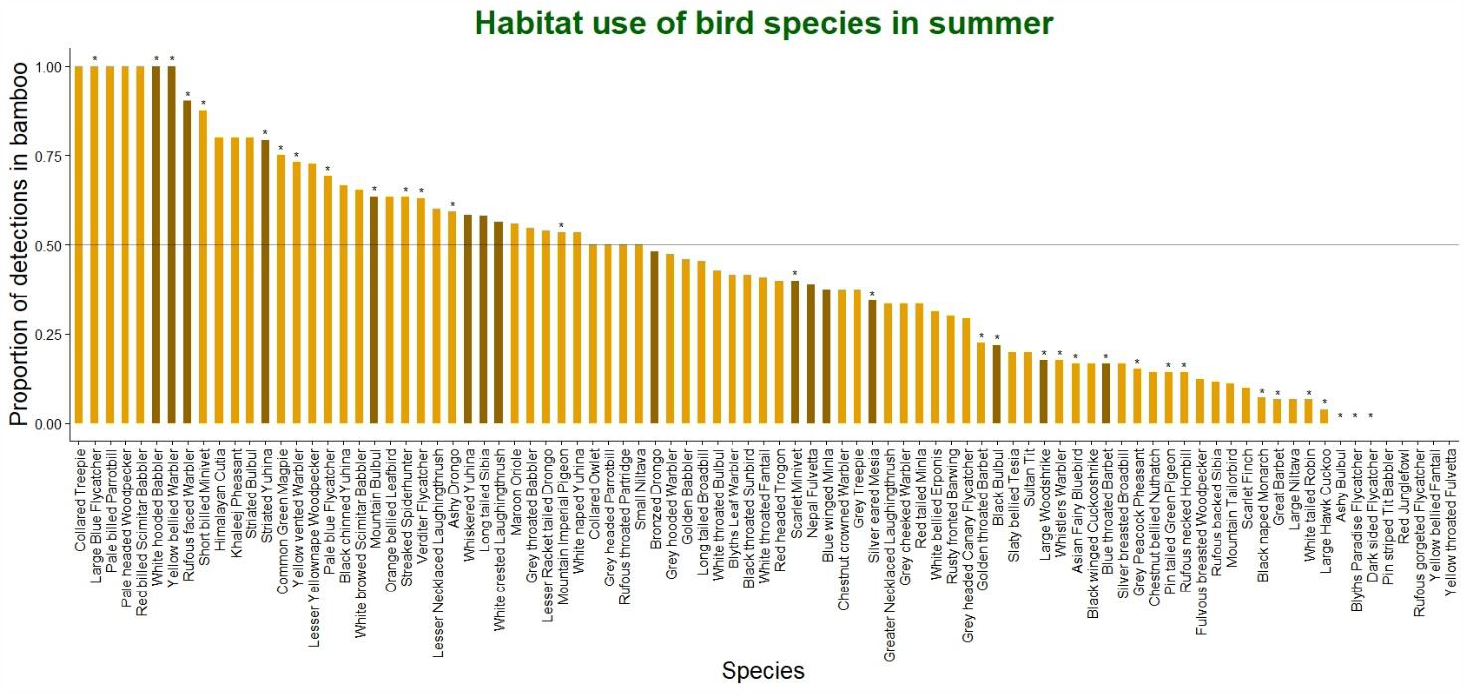
Habitat use scores of bird species in summer. Higher values indicate disproportionately high use of bamboo and lower indicate greater use of rainforest. Species are arranged in descending order of these values. Darker bars represent species that contribute to a cumulative 50% dissimilarity between the two habitats. Asterisk (*) above the bars indicate a significant contribution to the observed dissimilarity.

### Bird community composition

The Bray-Curtis index on transformed counts indicated that each habitat and season harboured a distinct bird community. The first NMDS axis separated bird communities in bamboo and rainforest habitats while the second axis separated winter and summer seasons (stress = 0.16, Fig. 5). PERMANOVA results indicated that the bird communities across habitats and seasons were significantly different (*R*^*2*^ = 0.32, *F*_3,24_ = 3.75, *p* < 0.01).

**Figure 5.**
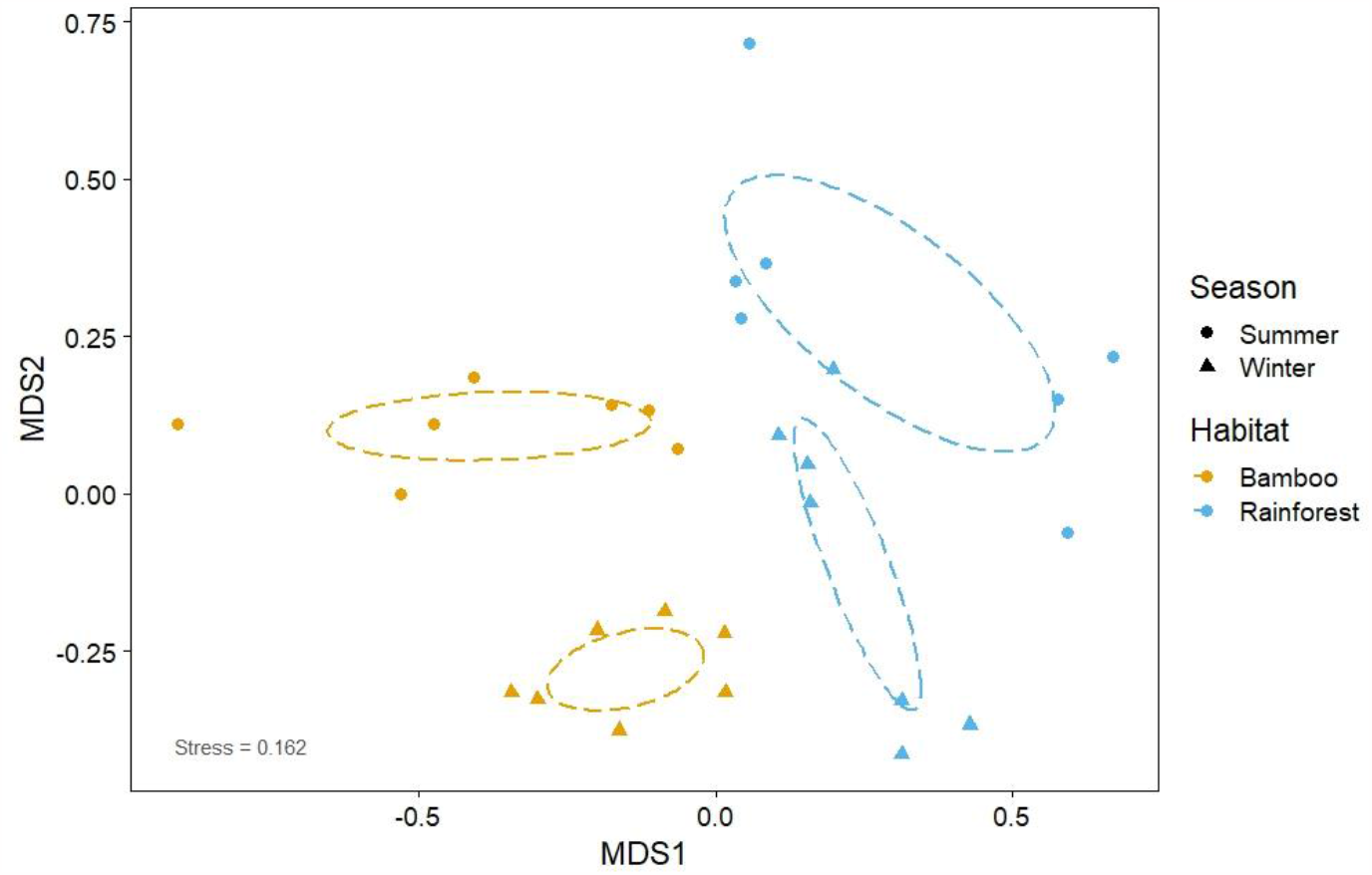
Non-metric dimensional scaling representing dissimilarity between winter and summer bird communities in bamboo and rainforest based on the Bray-Curtis index on Hellinger-transformed count data. Points represent transects in each season and habitat. Ellipsoids are centred around the mean and represent 95% confidence intervals around the points.

The similarity percentage scores revealed the most important species contributing to the overall dissimilarity between the bird communities in bamboo and rainforest in each season. In winter, these were mostly species with lower values of habitat-use scores, (i.e., species that were using the rainforest habitat more). These were the Red-tailed Minla (*Minla ignotincta*), Rusty-fronted Barwing (*Actinodura egertoni*), Blue-winged Minla (*Actinodura cyanouroptera*) and Rufous-vented Yuhina (*Yuhina occipitalis*). Some species with higher habitat-use values also had high similarity percentage scores, for instance, the White-hooded Babbler and Yellow-bellied Warbler. Other species with high contribution to the dissimilarity in winter bird communities included generalist species such as the Whiskered Yuhina (*Yuhina flavicollis*; dissimilarity score = 0.068), Nepal Fulvetta (*Alcippe nipalensis*; 0.037), White-crested Laughingthrush (*Garrulax leucolophus*; 0.018) and the Long-tailed Sibia (*Heterophasia picaoides*; 0.016). Only the Red-tailed Minla and the Rusty-fronted Barwing had significant scores in winter (Figure 3; Table A1, Supplementary data).

In summer, the most important species contributing to the dissimilarities were mostly species with higher scores of habitat use (i.e., species that were using bamboo more) such as the Striated Yuhina (*Staphida castaniceps*), White-hooded Babbler, Yellow-bellied Warbler and Rufous-faced Warbler (*Abroscopus albogularis*). Species with low habitat-use scores (more rainforest use) included the Blue-throated Barbet (*Psilopogon asiaticus*), Large Woodshrike (*Tephrodornis virgatus*) and Himalayan Black Bulbul (*Hypsipetes leucocephalus*). Other species that had high dissimilarity scores included generalists, like the Silver-eared Mesia (dissimilarity score = 0.03; *Leiothrix argentauris*), Nepal Fulvetta (0.03) and the Whiskered Yuhina (0.029, Figure 4; Table A1, Supplementary data).

### Arthropod community composition

Sampling from both seasons and habitats yielded 7,354 individuals representing 21 orders in total. The major orders were – Araneae, Hymenoptera (> 90% of hymenopterans were represented by ants), Acarina, Diptera, Blattodea, Hemiptera, Orthoptera, Psocoptera, Coleoptera and Collembola. Estimated jackknife order level richness was not particularly different in bamboo and rainforest across both seasons (Figure 6).

**Figure 6.**
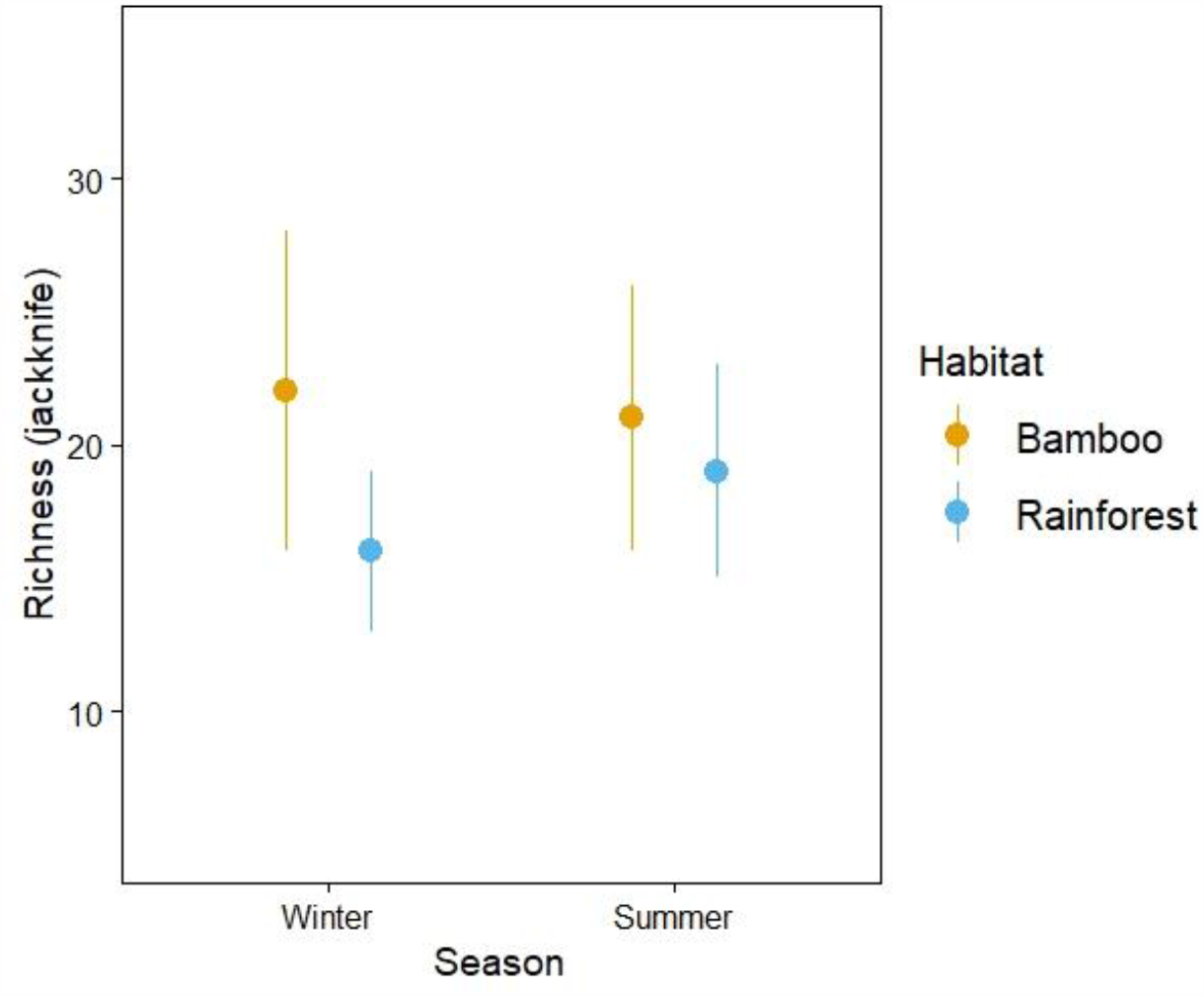
Estimated jackknife order richness of arthropods across habitats and seasons. Error bars represent 95% confidence intervals.

Arthropod abundances were higher in summer than in winter, across habitats and methods (Fig. A1, A2; Supplementary data) but within a habitat and across seasons, patterns were not discernible due to the high abundances of Hymenoptera and Isoptera in both habitats in the summer. When Hymenoptera and Isoptera were excluded, rainforest showed slightly higher mean abundances in the summer across habitats, with values from winter being largely similar (Fig. A2, Supplementary data). Other orders showed differences in their abundances between particular habitats, seasons and methods. For instance, Blattodea and Collembola were almost entirely absent in foliage but were fairly abundant in pitfall traps. Hemiptera, Araneae and Pscoptera seemed to show the opposite trend, with more abundance in bagging than pitfall traps (Fig. A3, A4, Supplementary data). Abundances of Hemiptera, Coleoptera, Collembola and Araneae did not vary considerably between habitats in each season, irrespective of the method (excluding sheaths). The mean number of arthropods per point in the sheaths in bamboo seemed to be similar to the mean values from the other methods and habitats in winter (Fig. A5, Supplementary data). Order-wise abundances in sheaths revealed that some orders such as Coleoptera, Hemiptera and Araneae were more abundant than others.

The arthropod community exhibited significant differences across habitats and seasons (*R*^*2*^ = 0.09, *F*_3,234_ = 8.04, *p* < 0.01), albeit the stress was on the higher side (stress = 0.19, Figure 7). The first NMDS axis separated arthropod communities across seasons and the second axis divided the arthropod communities across habitats, indicating the presence of distinct arthropod communities. Further, we found that winter arthropod communities across bamboo substrates were significantly different; the foliage-dwelling arthropod community in bamboo separated well from the culm sheath community, in the first NMDS axis (stress = 0.20; *R*^*2*^ = 0.06, *F*_1,58_ = 0.81, p < 0.01; Figure 8).

**Figure 7.**
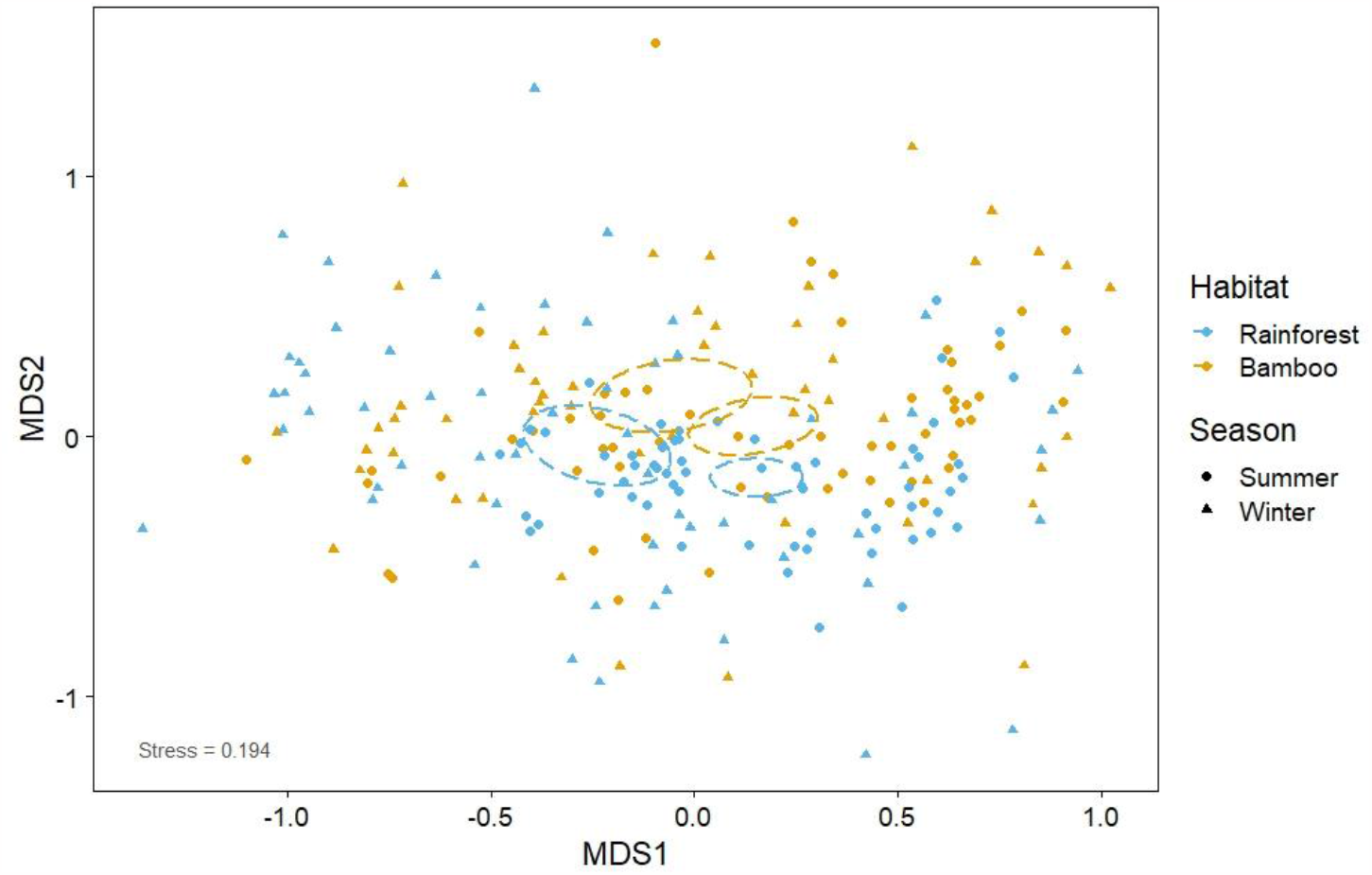
Non-metric dimensional scaling representing dissimilarity between winter and summer arthropod communities in bamboo and rainforest based on the Bray-Curtis index on Hellinger transformed count data. The points represent arthropod samples within season and habitat. Ellipsoids are centred around the mean and represent 95% confidence intervals around the points.

**Figure 8.**
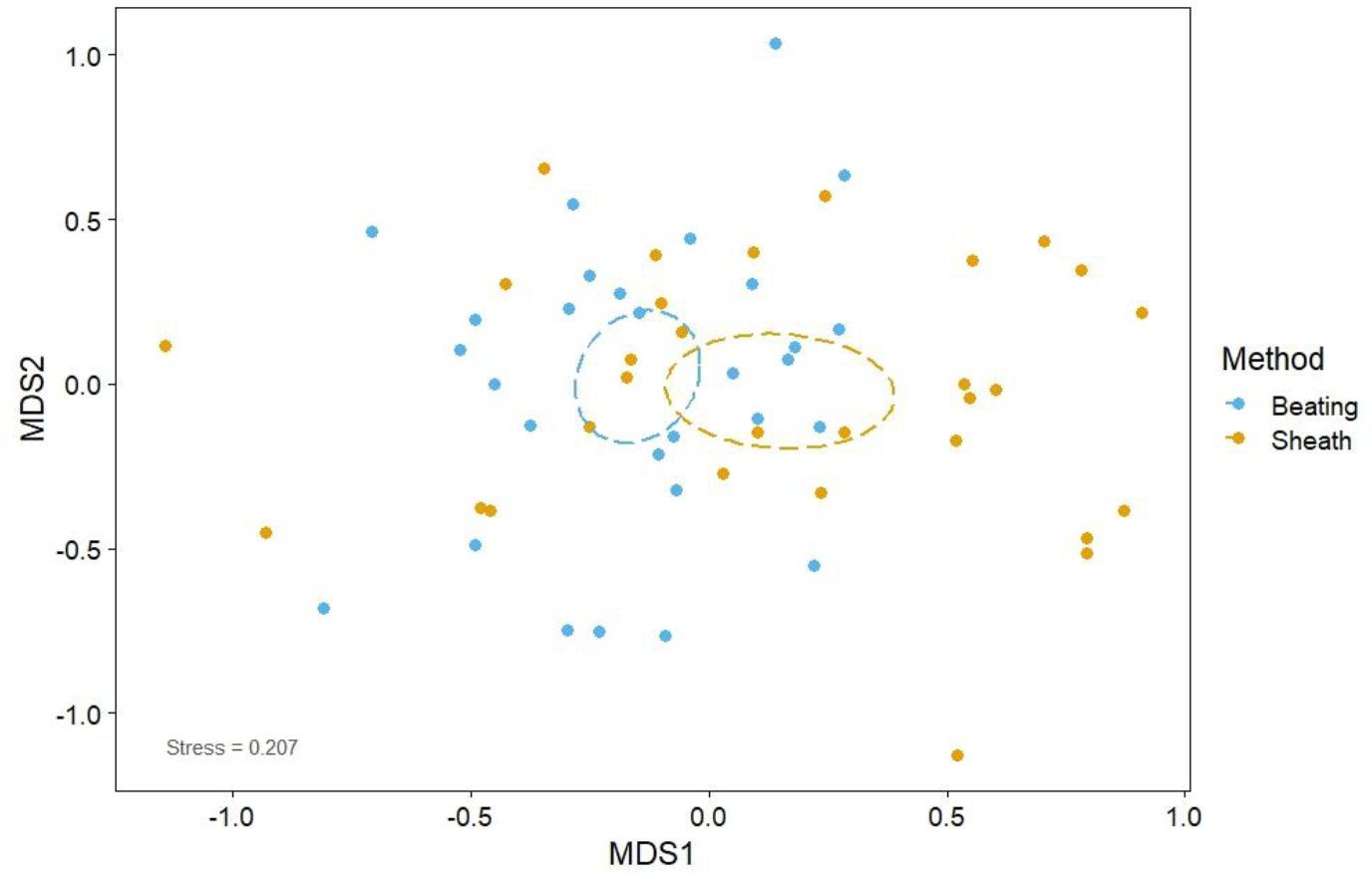
Non-metric dimensional scaling representing dissimilarity between arthropod communities within different substrates in bamboo based on the Bray-Curtis index on Hellinger transformed count data. The points represent random points used for sampling arthropods within season and habitat. Ellipsoids are centred around the mean and represent 95% confidence intervals around the points

## Discussion

### Bird communities

This study provides the first clear evidence of habitat specialisation in bamboo by Eastern Himalayan birds. The species that specialise on bamboo were the White-hooded Babbler, Pale-billed Parrotbill, Red-billed Scimitar Babbler, Yellow-bellied Warbler and Pale-headed Woodpecker. The habitat-use scores of these species were either one or very close to one in both seasons, ruling out a seasonal specialisation on bamboo. Estimated jackknife species richness did not show considerable differences between bamboo and rainforest in both seasons, contradicting patterns observed in Neotropical studies, where bamboo has higher bird species richness (Reid et al. 2004, Rother et al. 2013). Community composition was distinct across seasons and habitats (Figure 5), indicating that bamboo harbours distinct bird communities that are similar in species richness when compared with rainforest – indicating the importance of bamboo in supporting biodiversity in this region.

We found that the bamboo bird community was mostly comprised of the bamboo specialist species in both seasons along with other generalist species occurring at this elevation. Examples of generalists include the Nepal Fulvetta, Golden Babbler (*Cyanoderma chrysaeum*), Whiskered Yuhina, Black-throated Sunbird (*Aethopyga saturate*) and White-throated Fantail (*Rhipidura albicollis*) (habitat-use values close to or equal to 0.5, Figure 3, Figure 4). Frugivores, such as hornbills, barbets and pigeons were noticeably absent in bamboo but occurred in rainforest. Bamboo perhaps does not offer suitable resources for frugivores, thereby explaining their absence (Rother et al. 2013). Other species that were absent included bark-feeders such as the two nuthatch species, Rufous-backed Sibia (*Leioptila annectens*) and the Fulvous-breasted Woodpecker (*Dendrocopos macei)*, and a few understorey insectivores such as the Yellow-throated Fulvetta, Pin-striped Tit-Babbler and White-tailed Robin (*Myiomela leucura*). These species may require specific microhabitats or resources that are absent in bamboo and hence might avoid using bamboo. Habitat-use scores were perhaps misleading for some species such as the Mountain Imperial Pigeon (*Ducula badia*), Great Barbet (*Psilopogon virens*), Himalayan Cutia (*Cutia nipalensis*) and Yellow-browed Warbler (*Phylloscopus inornatus*), as they were detected on the few large trees within the bamboo habitat but rarely on bamboo itself. These results could mean that while bamboo might be important for specialists, it is not unattractive to generalists either, concurring with (Rother et al. 2013), who found a considerable number of non-bamboo specialists using bamboo habitats in the Neotropics.

### Arthropod communities

A key finding was that arthropod communities in bamboo and rainforest were distinct, even across seasons (Figure 7). Because order-level richness of arthropods did not differ considerably between habitats and seasons, variation in abundances across orders is likely to be driving differences in community composition. Arthropod abundances were not dissimilar between habitats and within a season, after filtering out Hymenoptera and Isoptera (Fig. A3, A4, A5, Supplementary data). This is in contrast to patterns in the Neotropics, where arthropod abundances are much higher in bamboo than in forests, possibly because of higher biomass in bamboo (Reid et al. 2004). Though we expected a seasonal increase in abundance (Ghosh et al. 2011), the magnitude of the increase was unexpected, especially in Hymenoptera, whose abundances soared in summer in bamboo pitfall traps (Fig. A1, A4, Supplementary data). Ants seem to particularly favour bamboo in the summer, possibly for using it as a substrate for nesting (Klein et al. 1993, Schellerich-Kaaden et al. 1997, Davidson et al. 2006, Leite et al. 2013, Arruda et al. 2016). In addition, the abundances of orders such as Hymenoptera, Diptera, Blattodea and Collembola from pitfall traps seemed to be much higher than that of the branch beats, especially in summer in bamboo, while orders such as Hemiptera and Araneae showed the opposite trend (Fig. A3, A4, Supplementary data).

We found that the arthropod community in bamboo culm sheaths in winter was distinct from the bamboo foliage-dwelling arthropods, with arthropods in sheaths separating well even at the order level (Figure 8). This difference is not surprising, given that many other arthropod groups are also known to specialise on different substrates of bamboo (Louton et al. 1996, Hancock and Drew 1999, Kovac 2000, Campos 2013). Furthermore, only a few orders were abundant in the sheaths when compared with bamboo foliage (Fig. A5, Supplementary data). These differences further contribute to the contrast in community composition between the substrates and point to culm arthropods and possibly other bamboo substrates, being specialists themselves.

### Foraging behaviour

Though not the main focus of this study, our observations in the field indicate that only bamboo specialist species (barring the Yellow-bellied Warbler and the Collared Treepie), seek arthropods from within the culm sheaths in winter, while generalist birds inhabiting bamboo did not employ this method to forage. The Red-billed Scimitar Babbler and the White-hooded Babbler probe the culm sheaths and the internodes for arthropods using their long and hooked bills, while the parrotbills tear open the culm sheaths with their strong bills and also inspect the internodes. The culm sheaths are shed in summer, but the internodes were probed irrespective of the season. This behaviour is not unlike the foraging behaviour of bamboo insectivore specialists from the Neotropics, as they are also known to probe the internodes, open the bamboo culm sheaths and inspect existing holes in the culms to hunt for arthropods (Hilty et al. 1979, Rodrigues et al. 1994, Parker et al. 1997, Areta and Cockle 2012, Leite et al. 2013). These strategies are likely to make these Eastern Himalayan bird species specialists at foraging for arthropods in bamboo, enabling them to access an entire community of arthropods that is perhaps unavailable to other birds (Figure 8). This is especially the case during winter when arthropod abundances are the lowest and consequently, competition for arthropod prey is the highest in the Himalaya (Ghosh et al. 2011, Srinivasan et al. 2018). Specialisation in bamboo might have evolved to allow birds to evade competition during periods of resource scarcity in winter and help in providing a consistent and reliable source of food. Newer techniques such as fecal DNA metabarcoding should be used to compare the diets of bamboo and rainforest species in conjunction with arthropod data (Stillman et al. 2022).

### Alternative hypotheses, further questions and conservation

A main caveat of our study is that we explore only one possible reason for the association of birds with bamboo – dietary preferences. There may be other mechanisms that might act in conjunction or even exclusively to diet, such as the structure and the composition of bamboo.

Bird associations with bamboo in the Neotropics are mainly with one genus of bamboo, *Guadua* (Kratter 1997, Lebbin 2007). The bamboo stands in our study were dominated by only one species of bamboo (*Dendrocalamus longispathus*) and while there could be a certain propensity for bamboo specialists to associate with the species in this genus, future studies comparing bird communities in different species of bamboo might help discover the affinities of bird species with other bamboo species in this region.

Different species of bamboo are inherently structured differently. Vegetation structure can play an important role in determining various life-history strategies and ecologies of species. For example, in addition to affecting overall resource availability, vegetation density and complexity can influence prey visibility, foraging success, predation risk, nest and roost sites, and consequently, habitat selection by birds (Whelan 2001, Raman 2006, Lebbin 2007, Socolar et al. 2013, Menon et al. 2019). The vegetation structure of bamboo could help in creating a less complex habitat than rainforest, as bamboo contains more homogenous leaf structures, providing a consistent and simpler search image for birds. This can help them forage more effectively and exploit a particular resource, facilitating habitat specialisation. This mechanism is hypothesised to be at play in the Neotropics, especially with bamboo specialist sallying flycatchers (Kratter 1997, Lebbin 2007, Socolar et al. 2013). While no sallying flycatchers are known to be exclusive to bamboo in the Eastern Himalaya, this mechanism could help explain the consistent foraging behaviour exhibited by the bamboo specialists in this region. Additionally, while the structure of bamboo might be less complex than that of rainforest, bamboo has denser foliage. This could function as potential cover for birds from predators, as the denser habitat might hamper larger raptorial species from hunting effectively (Reid et al. 2004). Call playback experiments can help test the predator refuge hypothesis in future studies.

Moreover, other factors such as patch size, patch area, patch shape, distance to adjacent bamboo patches and the quality of the surrounding matrix may prove to be vital for bamboo specialists to colonise and persist in bamboo stands. Some Neotropical bamboo specialists require large stands of bamboo and smaller patches harbour a subset of the community found in larger stands (Lebbin 2007, 2013). Further, the territory sizes of several bamboo specialists seemed to be smaller than non-bamboo specialists in the Amazon (Kratter 1997). Estimating home range sizes and the density of specialists in various patch sizes of different species of Eastern Himalayan bamboo might be a key area of research for future studies.

A vital question is, what happens to the inhabitants of bamboo when bamboo dies after mast seeding events? The ability of a species to disperse could be key to their survival, especially in a highly spatially and temporally fluctuating habitat such as bamboo. As a corollary, this could mean that birds that are strongly associated with bamboo could be vagile species, being able to successfully disperse to other bamboo habitats to ensure survival (Areta and Cockle 2012). This raises some interesting questions – are the Eastern Himalayan bamboo specialists highly vagile? What is the minimum distance required between bamboo patches for effective colonisation? Do the bird populations in scattered bamboo patches constitute a larger meta-community? Moreover, a few species also seem to specialise on the seeds from the mast seeding events in the Neotropics (Areta et al. 2009, Lees et al. 2021). Future studies can also investigate the existence of such species from the Eastern Himalaya.

Another important aspect to consider is climate change. Bamboo specialists might be considerably more vulnerable to the effects of climate change than generalists. If specialization is facultative, these species might be able to track temperature upslope and colonise bamboo stands at higher elevations; however, if suitable habitat does not exist at higher elevations or if they are unable to move upslope, local extirpation might be likely. Moreover, bamboo is sensitive to climate change and is likely to be adversely affected by warming temperatures (Srivastava et al. 2019). Other faunas strictly associated with bamboo are already threatened by the effects of climate change (Eronen et al. 2017).

Finally, bamboo is an important resource for resident communities because of its ubiquity and versatility (Bystriakova 2003). Therefore, an inherently fluctuating habitat is likely being made even more variable by human use. This could make some insectivorous birds, a group that is vulnerable to land-use change due to its habitat specificity, even more threatened (Sekercioglu et al. 2007). Habitat specialist birds are known to be the most vulnerable to anthropological change, as specialists are being replaced by generalists worldwide (Clavel et al. 2011). This could place bamboo specialist insectivorous species in this region at particular risk from global change drivers.

## Conclusions

Our study is the first that systematically documents and confirms the existence of bamboo specialists in the Eastern Himalaya. Bird communities in bamboo and rainforest show considerable differences, as do arthropod communities between the two habitats and across seasons. Many generalist bird species were also found to be using bamboo habitats; therefore, bamboo provides habitat to non-bamboo specialists as well. Arthropod communities in/on different substrates in bamboo were distinct. The unique foraging behaviour of bamboo specialist bird species combined with their dietary preferences can potentially explain their association with this habitat. We finally discuss alternative hypotheses, strategies and future questions that might help explain the unique ecologies of species in bamboo and might help inform the conservation of these species and their habitats.

## Acknowledgements

This study was carried out as a part of SS’s Master’s dissertation. We thank the National Centre for Biological Sciences-Tata Institute of Fundamental Research (NCBS-TIFR) and the Nature Conservation Foundation for institutional and administrative support. We are also thankful to the Arunachal Pradesh Forest Department for issuing permits to carry out this study (permit no. CWL/GEN/2018-19/Pt.IX/NG/3480-81). SS is grateful to The Habitats Trust for providing him with a grant for this study. Hari Sridhar and T. R. Shankar Raman provided useful discussions. Rinchen Angmo and Yeswanth H.M. helped greatly with arthropod identification. We also thank Mangal Rai, Nima Tamang, S.K. Pradhan, Bharat Tamang and Binod Munda for all their help and support during fieldwork.

## Author Contributions

SS and U.S. conceived and designed the study. AB, DKS, SR, and SS collected the data. SS analysed the data and was supervised by US. SS wrote the original draft and US and SS contributed in refining subsequent drafts.

