## Supplementary data for "Poles apart: the structure and composition of the bird community in bamboo in the Eastern Himalaya"

Supplementary Material/Appendix

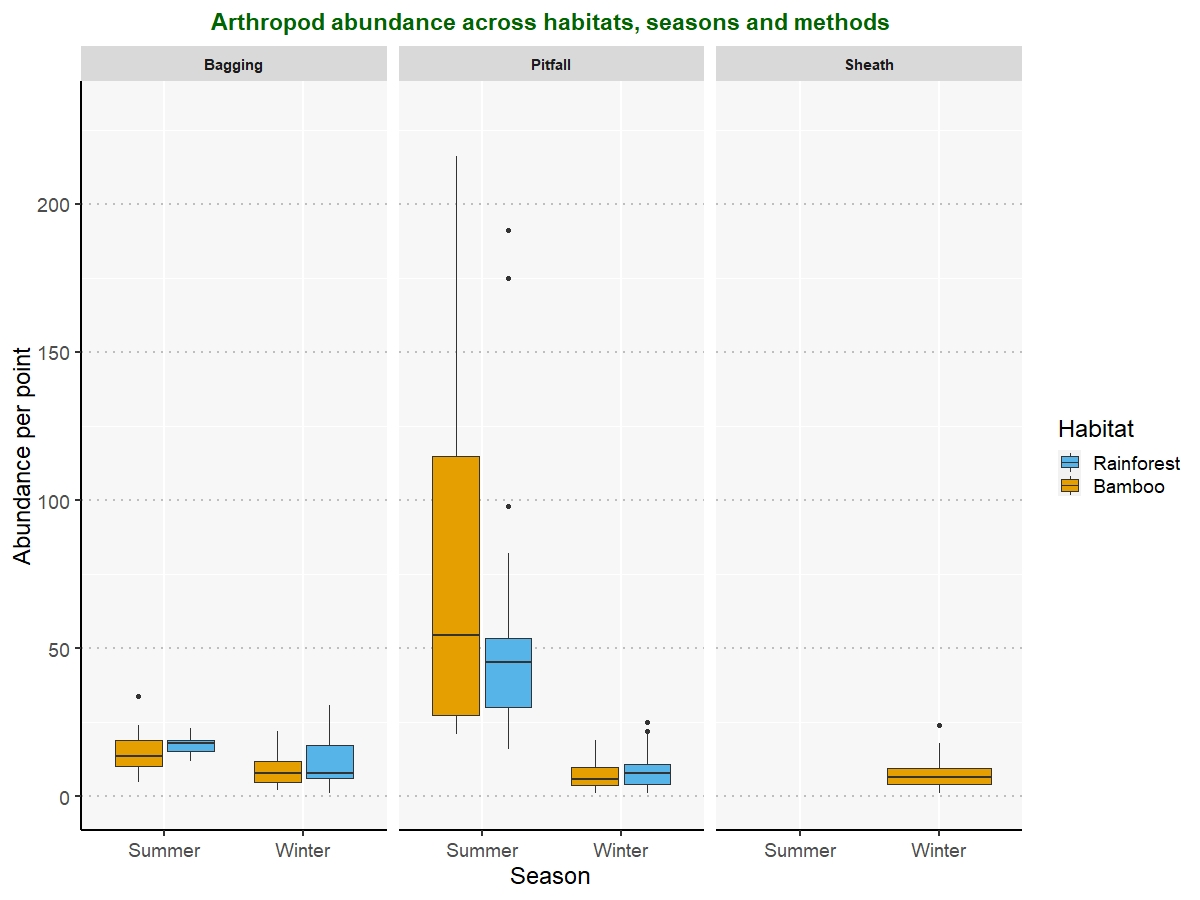

**Figure A1** - Arthropod abundance per-point across seasons and habitats. The whiskers represent the 1.5 times the inter-quartile range. The points are outliers.

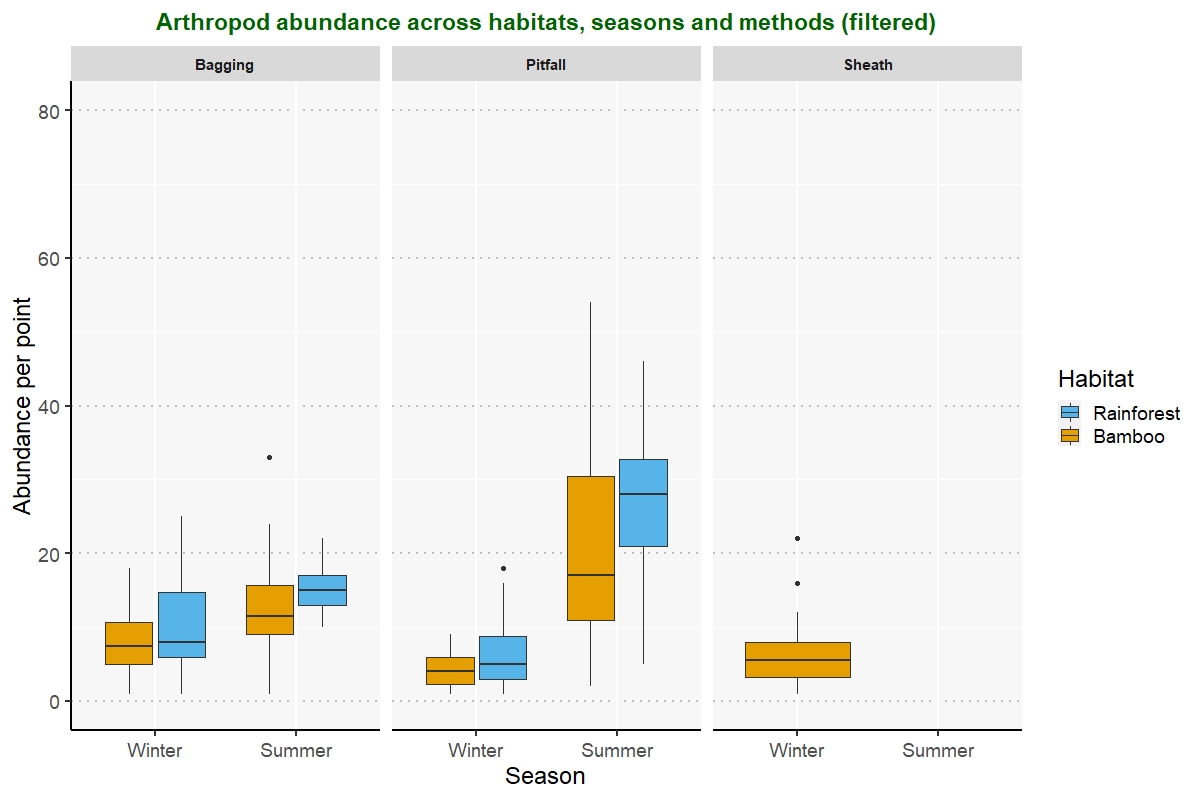

**Figure A2** – Per point arthropod abundance after filtering out Hymenoptera and Isoptera. The whiskers represent the 1.5 times the inter-quartile range. The points are outliers.

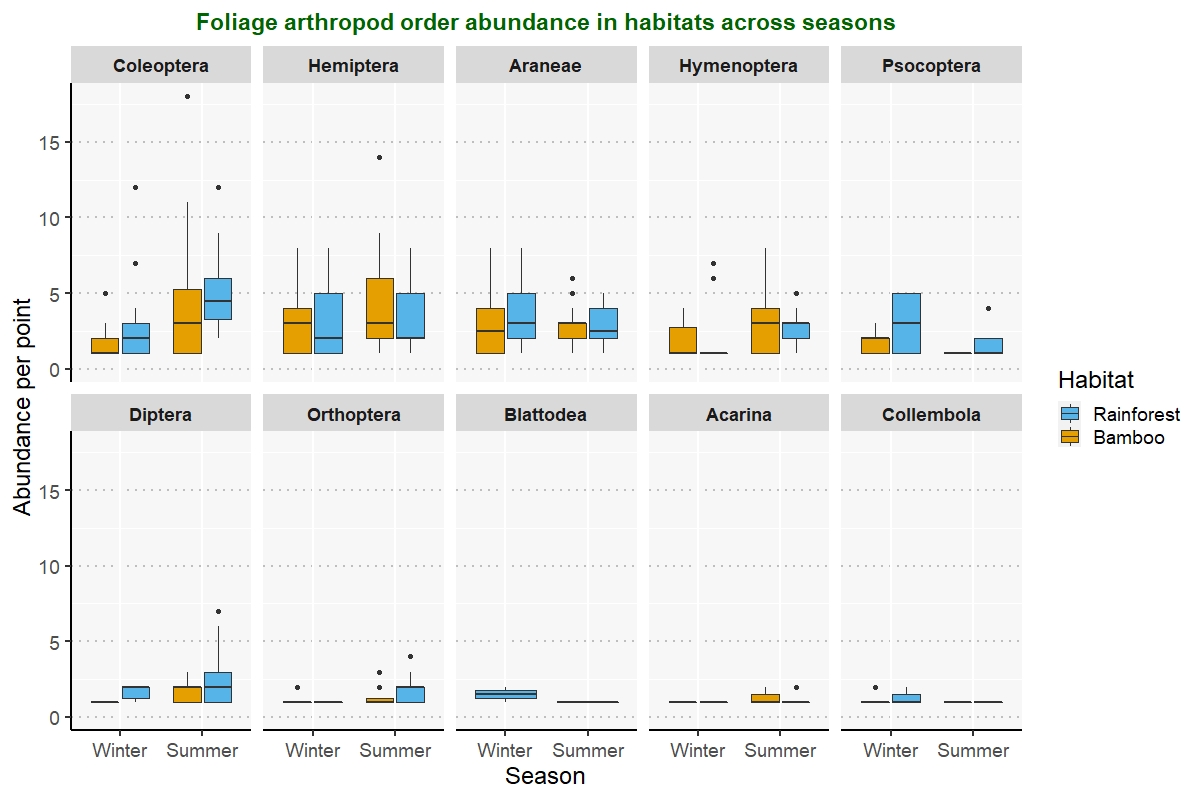

**Figure A3** – Per point foliage arthropod abundance of the 10 most abundant orders in both habitats and seasons. The whiskers represent the 1.5 times the inter-quartile range. The points are outliers.

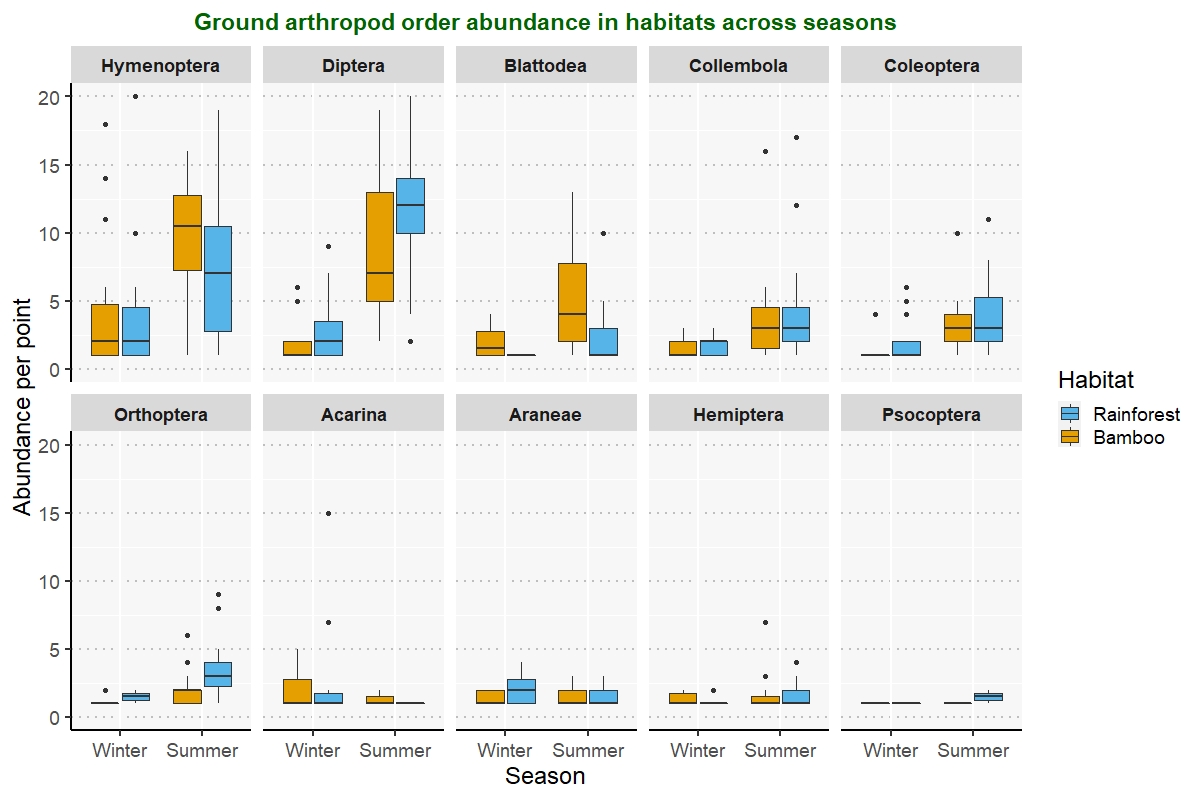

**Figure A4** – Ground arthropod order abundance per point, across seasons and methods. Only the 10 most abundant families are represented. The whiskers represent the 1.5 times the inter-quartile range. The points are outliers.

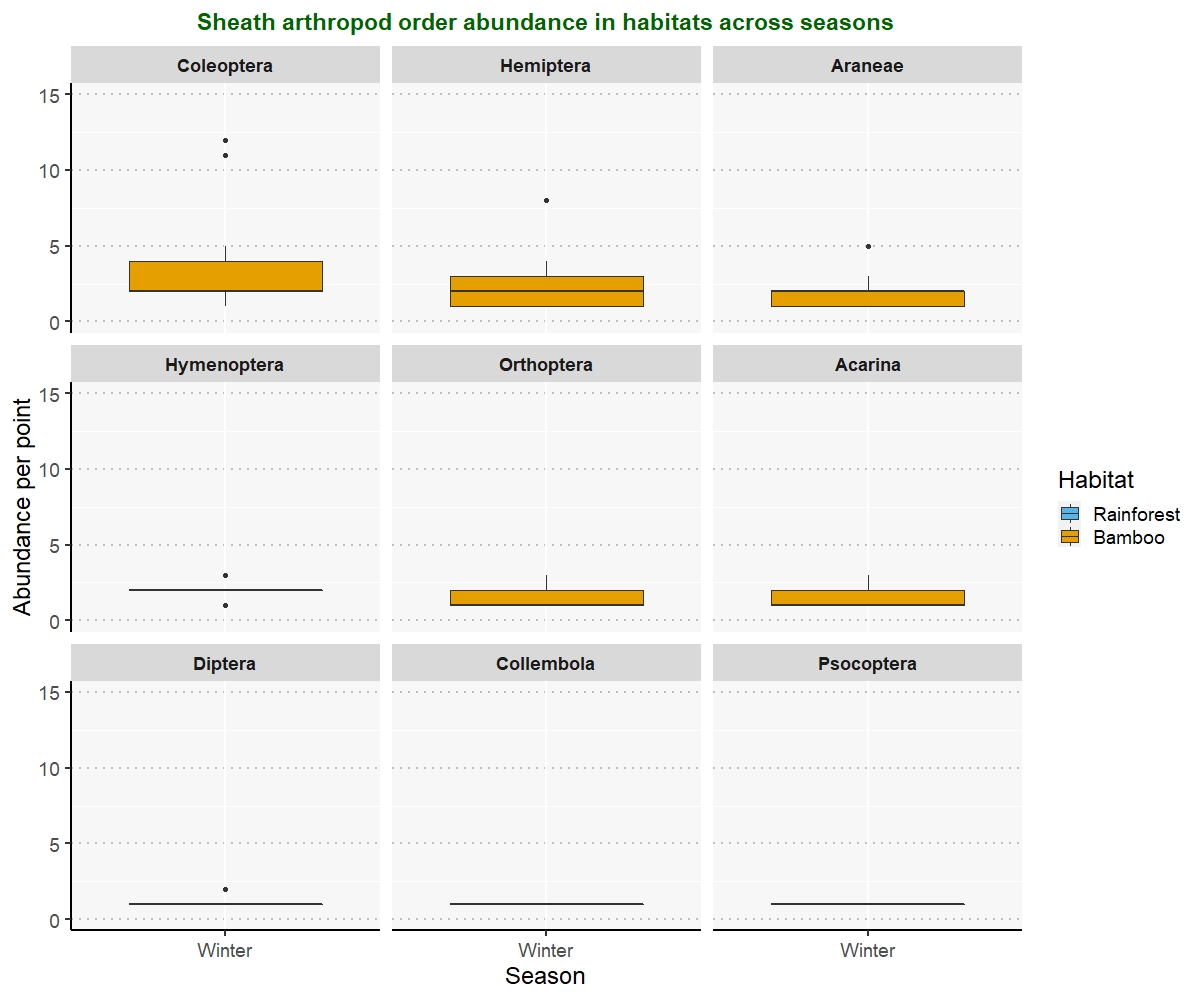

**Figure A5** – Per point sheath arthropod abundance of the 10 most abundant orders in bamboo in winter. Sheaths were only sampled in winter. The whiskers represent the 1.5 times the inter-quartile range. The points are outliers.
